# Liquefaction of the brain following stroke shares a similar molecular and morphological profile with atherosclerosis and mediates secondary neurodegeneration in an osteopontin dependent mechanism

**DOI:** 10.1101/264275

**Authors:** Amanda G. Chung, Jennifer B. Frye, Jacob C. Zbesko, Eleni Constantopoulos, Megan Hayes, Anna G. Figueroa, Danielle A. Becktel, W. Antony Day, John P. Konhilas, Brian S. McKay, Thuy-Vi V. Nguyen, Kristian P. Doyle

**Affiliations:** Department of Immunobiology, University of Arizona, Tucson, Arizona, 85719. USA; Department of Physiology and Sarver Molecular Cardiovascular Research Program, University of Arizona, Tucson, Arizona, 85724. USA; Department of Ophthalmology and Vision Science, University of Arizona, Tucson, Arizona, 85719. USA; Arizona Health Sciences Center Imaging Core Facility, Arizona Research Labs, University of Arizona, Tucson, AZ, 85719. USA; Department of Neurology, University of Arizona, Tucson, Arizona, 85719. USA; Arizona Center on Aging, University of Arizona, Tucson, Arizona, 85719. USA

## Abstract

Here we used mouse models of heart and brain ischemia to compare the inflammatory response to ischemia in the heart, a protein rich organ, to the inflammatory response to ischemia in the brain, a lipid rich organ. We report that ischemia-induced inflammation resolves between 1 and 4 weeks in the heart compared to between 8 and 24 weeks in the brain. Importantly, we discovered that a second burst of inflammation occurs in the brain between 4 and 8 weeks following ischemia, which coincided with the appearance of cholesterol crystals within the infarct. This second wave shares a similar cellular and molecular profile with atherosclerosis and is characterized by high levels of osteopontin (OPN) and matrix metalloproteinases (MMPs). In order to test the role of OPN in areas of liquefactive necrosis, OPN^-/-^ mice were subjected to brain ischemia. We found that at 7 weeks following stroke, the expression of pro-inflammatory proteins and MMPs was profoundly reduced in the infarct of the OPN^-/-^ mice, although the number of cholesterol crystals was increased. OPN^-/-^ mice exhibited faster recovery of motor function and a higher number of neuronal nuclei (NeuN) positive cells in the peri-infarct area at 7 weeks following stroke. Based on these findings we propose that the brain liquefies after stroke because phagocytic cells in the infarct are unable to efficiently clear cholesterol rich myelin debris, and that this leads to the perpetuation of an OPN-dependent inflammatory response characterized by high levels of degradative enzymes.

**Significance Statement:** The inflammatory response to ischemia in the brain is different to the response to ischemic injury in other organs. In the brain, and for unknown reasons, dead tissue liquefies in response to ischemia by the process of liquefactive necrosis. However, the data we present here demonstrate that there is overlap between the pathophysiology of liquefactive necrosis and atherosclerosis. Specifically, we show that chronic stroke infarcts contain foamy macrophages, cholesterol crystals, high levels of OPN and MMPs, and a similar cytokine profile to atherosclerosis. Therefore, because cholesterol is a central component of myelin, liquefactive necrosis in response to stroke may be caused by an inflammatory response to cholesterol-rich myelin debris that is driven in large part by OPN and MMPs.

## Introduction

In response to ischemia, the brain degenerates by the process of liquefactive necrosis while the heart degenerates by coagulative necrosis (Robbins et al., 2010). Liquefactive necrosis is a chronic inflammatory response that persists for months following a stroke and occurs for unknown reasons (Doyle et al., 2015; Nguyen et al., 2016; Zbesko et al., 2018). Cholesterol is a central structural component of myelin. This means that in the aftermath of a brain injury, the immune system must clear away substantially more cholesterol than in the aftermath of an injury to an alternative area of the body such as the heart following a myocardial infarction. Therefore, the goal of this research was to understand if the brain undergoes liquefactive necrosis in response to stroke due to the high cholesterol content of the brain.

The rationale for this is that in cardiovascular disease, atherosclerotic plaques develop within arteries when macrophages ingest excessive cholesterol (Moore et al., 2013). Macrophages slowly develop into large proinflammatory foam cells that contain numerous lipid-laden internal cytoplasmic vesicles. Foam cell formation is dictated by the balance between extracellular lipid uptake and reverse lipid transport from the intracellular compartment to extracellular lipid acceptors, such as high-density lipoprotein (HDL). When this balance tilts toward excessive cholesterol, it leads to cell death and chronic inflammation through multiple mechanisms. An example of one such mechanism is that free cholesterol is toxic to cells. Consequently, it must be shuttled to the endoplasmic reticulum (ER) to be esterified and stored in lipid droplets, or effluxed from the cell via transporters such as the ATP-binding cassette transporter, ABCA1 (Remmerie and Scott, 2018). When there is too much free cholesterol in the ER the esterification machinery can become overwhelmed leading to the accumulation of free cholesterol, toll-like receptor (TLR) signaling, and nuclear factor kappa-light-chain enhancer (NF-kB) activation (Remmerie and Scott, 2018). This inflammatory signaling, combined with ER stress and the accumulation of free cholesterol, can cause cell death by apoptosis (Remmerie and Scott, 2018). Apoptotic macrophages must be cleared by neighboring macrophages which may themselves have stressed lipid metabolism and an impaired ability to perform this clearance function. This can result in prolonged damage-associated molecular pattern (DAMP) mediated TLR signaling, chronic inflammation, and the further propagation of cell death (Remmerie and Scott, 2018).

Another problem associated with excessive cholesterol in a biological system is that the cholesterol transporter low density lipoprotein (LDL) can be oxidized. Oxidized LDL is a ligand for scavenger receptors and TLR4 and can thereby directly trigger pro-inflammatory signaling pathways (Remmerie and Scott, 2018). Furthermore, when cholesterol accumulation within a cell exceeds the solubility limit of the cell, cholesterol crystals form (Duewell et al., 2010). Cholesterol crystals cause physical damage to organelles by destabilization of the lysosomal compartment, rupture plasma membranes, and activate nucleotide-binding domain and leucine-rich repeat containing protein 3 (NLRP3) inflammasome signaling (Abela and Aziz, 2005; Duewell et al., 2010; Moore et al., 2013; Corr et al., 2016).

Consequently, the goal of this study was to investigate if the overloading of cholesterol by macrophages is not only involved in the pathophysiology of cardiovascular disease, but due to the fact that the brain is the most cholesterol-rich organ, containing approximately 20% of the body’s total cholesterol (Orth and Bellosta, 2012), if it is also a potential explanation for why brain tissue undergoes liquefactive necrosis following stroke. In support of this hypothesis, Cantuti-Castelvetri and colleagues recently demonstrated that in a toxin-induced model of demyelination, myelin debris can overwhelm the efflux capacity of microglia, resulting in the formation of cholesterol crystals and a chronic inflammatory response in aged mice (Cantuti-Castelvetri et al., 2018). Additionally, transcriptional profiling of macrophages at 7 days after spinal cord injury reveals that they closely resemble foam cells, with lipid catabolism representing their main biological process (Zhu et al., 2017).

Therefore, we used mouse models of myocardial infarction (MI) and ischemic stroke to compare the kinetics and characteristics of the inflammatory response to ischemia in each organ. We show that the brain takes substantially longer than the heart to heal, with the induction of a second wave of inflammation that coincides with the formation of cholesterol crystals within the infarct in the weeks following stroke. The second wave includes the expression of many of the same cytokines associated with atherosclerosis as well as high levels of OPN and MMPs. We then show that the chronic expression of these pro-inflammatory cytokines and proteolytic enzymes is reduced in OPN^-/-^ mice, and that these animals exhibit an accelerated recovery of motor function and a reduction in secondary neurodegeneration when compared to wildtype (WT) mice. These findings provide a potential explanation for why the brain liquefies following stroke, that is to say, the processing of cholesterol-rich myelin debris within phagocytic cells is overwhelmed, leading to the long-term OPN-dependent production of proteolytic enzymes.

## Materials and Methods

### Mice

Adult 10-12-week old male C57BL/6 (Stock No: 000664), BALB/c (Stock No: 000651), NLRP3^-/-^ (Stock No: 021302), CD36^-/-^ (Stock No: 019006), and OPN^-/-^ (Stock No: 025378) mice were purchased from the Jackson Laboratory (Bar Harbor, ME). All animals were housed under a 12-hour light/dark schedule with *ad libitum* access to food and water. All experiments were in accordance with protocols approved by the National Institutes of Health guidelines and the University of Arizona Institutional Animal Care and Use Committee.

### Stroke surgery

C57BL/6, NLRP3^-/-^, CD36^-/-^, and OPN^-/-^ mice underwent distal middle cerebral artery occlusion (DMCAO) + hypoxia (DH stroke), as described previously (Doyle et al., 2012). Briefly, mice were anesthetized by isoflurane inhalation, core body temperature was maintained at 37°C, and the skull exposed by creating an incision in the skin and temporalis muscle. The right middle cerebral artery (MCA) was identified and a microdrill was used to penetrate the skull to expose the underlying MCA. The meninges were cut and the vessel cauterized using a small vessel cauterizer (Bovie Medical Corporation). Surgical wounds were closed using Surgi-lock 2oc (Meridian Animal Health). Mice were then immediately transferred to a hypoxia chamber (Coy Laboratory Products) containing 9% oxygen and 91% nitrogen for 45 minutes. Hypoxia increases the size of the infarct and reduces infarct volume variability in C57BL/6 mice (Doyle et al., 2012). Sham surgeries were performed identically to stroke surgeries, except for cauterization of the DMCA. BALB/c mice underwent DMCAO without hypoxia. BALB/c mice following DMACO have the same size infarct following DMCAO as C57BL/6 mice following DMCAO + hypoxia (Doyle et al., 2012). All mice received one dose of buprenorphine (0.1 mg/kg; Henry Schein) via subcutaneous injection immediately prior to surgery. Cefazolin antibiotics (25 mg/kg; Sigma-Aldrich) were subcutaneously administered immediately following surgery, and slow-release buprenorphine (1 mg/kg; ZooPharm) was subcutaneously administered 24 hours following surgery.

### Myocardial infarction

C57BL/6 mice were placed in the supine position on a heating pad and a light shone onto the neck region for transesophageal illumination to enable the larynx to be visualized through the mouth opening. The tongue was retracted and a guide wire inserted into the trachea. A 20G intravenous (IV) catheter was then laced over the guide wire and advanced into the trachea but not past the trachea bifurcation. The intubation tubing was then connected to a rodent ventilator. A left thoracotomy was then performed by a left lateral incision and transection of the third rib using dissecting scissors. An 8-0 polyethylene suture was threaded underneath the left coronary artery, perpendicular to the long axis of the heart. The ligature was tied and blanching of the myocardium was visually verified. The thoracotomy was then closed in 3 layers: (1) intercostal ribs and muscle, (2) the pectoral muscle, and (3) the skin layers using 7-0 nylon absorbable suture. After each surgery was completed, the expiration line of the ventilator was briefly occluded in order to evacuate the pleural space and establish proper intrapleural pressure. Mice were extubated and recovered on a heating pad. Total time under anesthesia was less than 30 minutes.

### Tissue processing

C57BL/6 mice were euthanized at 24 hours, 1 week, 4 weeks, 7 weeks, 8 weeks, and 24 weeks after stroke by deep isoflurane (JD Medical) anesthesia followed by exsanguination and intracardial perfusion with 0.9% saline. BALB/c mice were euthanized at 7 weeks following stroke by the same method. NLRP3^-/-^ and CD36^-/-^ mice were euthanized at both 24 hours and 7 weeks, while OPN^-/-^ mice were euthanized at 24 hours, 1 week, and 7 weeks. Brains and hearts were extracted, and either (i) snap frozen in liquid nitrogen for biochemical assays, (ii) post-immersed in 4% paraformaldehyde (PFA) and transferred to 30% sucrose for histological and immuno staining processing, (iii) placed in 2.5% glutaraldehyde and 2% PFA for electron microscopy processing, or (iv) placed in 2,3,5-triphenyltetrazolium chloride (TTC) for infarct size measurement. For histological and immunostaining, frozen coronal sections (40 µm or 6 µm) were taken through the entire brain using a Microm HM 450 sliding microtome (Thermo Fisher Scientific).

### Immunostaining

Immunostaining was performed on free-floating brain and paraffin embedded heart sections using standard protocols. Primary antibodies against neuronal nuclei (NeuN, 1:500; Millipore Sigma, Cat. No. MAB377, RRID: AB2298772), CD3e (1:1000; BD Biosciences, Cat. No. 550277, RRID: AB393573), CD68 (1:1000; Bio-Rad, Cat. No. MCA1957GA RRID: AB324217), B220 (1:500; BD Biosciences, Cat. No. 553085, RRID: AB394615), osteopontin (OPN, 1:500; Abcam, Cat. No. AB218237, RRID: AB2732079), ionized calcium-binding adapter molecule 1 (IBA-1, 1:500; Wako Cat. No. 019-19741, RRID: AB839506) and glial fibrillary acidic protein (GFAP, 1:2000; Millipore Sigma, Cat. No. AB5541, RRID: AB177521) were used in conjunction with the appropriate secondary antibody and ABC Vector Elite (Vector Laboratories) and 3,3’-diaminobenzidine kits (Vector Laboratories) for visualization. Sections were imaged using a digital Keyence BZ-X700 light and fluorescent microscope.

### Fluoro-Jade C staining

Brain sections were mounted and dried on a slide warmer. Slides were incubated in a 1:10 solution of sodium hydroxide in 80% ethanol for 5 minutes. Slides were washed with 70% ethanol for 2 minutes followed by a wash with distilled water for 2 minutes. Slides were incubated for 10 minutes in a 1:10 solution of potassium permanganate (Sigma-Aldrich) in distilled water followed by a wash with distilled water for 2 minutes. Slides were incubated in Fluoro-Jade C (Biosensis) and DAPI (4’,6-Diamidino-2-Phenylindole, double stranding DNA staining; Thermo Fisher Scientific) for 10 minutes in the dark. Slides were then rinsed 3 times in distilled water for 1 minute per rinse, cleared in xylene, and coverslipped with Entellan (Electron Microscopy Sciences). Sections were imaged using a digital Keyence BZ-X700 light and fluorescent microscope.

### Nissl staining

Brain and heart sections were mounted and hydrated through descending concentrations of ethanol, and subsequently dipped in distilled water. Sections were then immersed in a 0.5% Nissl (Sigma-Aldrich) solution for 1 minute, rinsed in water, differentiated in a 0.25% acetic alcohol solution, and then dehydrated through ascending concentrations of ethanol. Finally, sections were defatted in xylene, and coverslipped with Entellan. Sections were imaged using a digital Keyence BZ-X700 light and fluorescent microscope.

### Trichrome staining

Trichrome staining was performed on both mounted brain and heart sections (n=3-4 mice/experimental group) to visualize collagen deposition in infarcted tissue. Staining was performed using a trichrome staining kit (abcam, Cat. No. ab150686), following the kit’s protocol. Briefly, Bouin’s fluid was preheated in a water bath to 56-64°C and tissues were incubated in Bouin’s fluid for 1 hour followed by 10 minutes of cooling. Sections were rinsed in tap water until clear and once in distilled water. Sections were then stained using Weigert’s Iron Hematoxylin for 5 minutes and rinsed in running tap water for 2 minutes. Sections were placed in Briebirch Scarlet/Acid Fuchin solution for 1 minute and subsequently rinsed in water. Sections were then differentiated in Phosphomolybdic/Phosphotungstic acid solution for 10-15 minutes followed by application of Aniline Blue solution for another 5-10 minutes, and finally rinsed in distilled water. Acetic acid solution (1%) was applied to the sections for 3-5 minutes followed by dehydration in 95% ethanol solution, 2 times in 100% ethanol, defatted in xylene, and coverslipped with Entellan. Sections were imaged using a digital Keyence BZ-X700 light and fluorescent microscope.

### Infarct evaluation

To visualize the region of infarction, animals were sacrificed at 24 hours post-stroke, and their brains processed for TTC staining (n=5 mice/experimental group). Brains were sectioned at 1 mm tissue thickness, and each section was immersed in 1.5% TTC in phosphate buffered saline (PBS) at 37^°^C for 15 minutes, and then fixed in 10% formalin. The area of the ipsilateral hemisphere and the area of the infarct on each section were measured using NIH ImageJ analysis software. Measurements were multiplied by the distance between sections (1 mm), and then summed over the entire brain to yield volume measurements.

### Multiplex immunoassay

Snap-frozen brain and heart tissue samples (n=5-11 mice/experimental group) were sonicated in ice-cold 0.1M PBS containing 1% triton-X and 0.1% sodium deoxycholate, Protease Inhibitor Cocktail (1:100; Millipore Sigma), and Phosphatase Inhibitor Cocktail 2 (1:100; Millipore Sigma). Following centrifugation, 13,000g at 4^°^C for 15 minutes, the total soluble protein concentration of each supernatant was measured using a Direct Detect Infrared Spectrometer (MilliporeSigma). Total MMP, OPN, and cytokines/chemokines concentrations were then measured using mouse multiplex magnetic bead kits (Milliplex Multiplex Assays, Millipore Sigma), according to the manufacturer’s protocols and recommendations. Each lysate sample, standard, and quality control were measured in duplicate. Plates were read using a MAGPIX instrument (Luminex), and results were analyzed using MILLIPLEX Analyst 5.1 software (Millipore Sigma).

### Electron microscopy

Brains (n=3 mice) used for transmission and scanning electron microscopy (EM) were fixed in 2.5% glutaraldehyde + 2% PFA in 0.1 M Pipes buffer for 1 hour at room temperature or overnight at 4^°^C. Following washes in 0.05 M Pipes buffer/0.05 M glycine and washes in 0.1 M Pipes buffer, dissected stroke regions from each brain were then post-immersed in 1% osmium tetroxide for 1 hour. After fixation, samples underwent washes in deionized water (DIW). Samples for transmission EM were then block stained in 2% aqueous uranyl acetate, washed in DIW, dehydrated through a graded series of ethanols, infiltrated with 1:1 alcohol/Spurr’s resin overnight, and embedded in 100% Spurr’s resin overnight at 60^°^C. Samples for scanning EM were dehydrated, incubated in hexamethyldisilazan, and then air dried. Sections for transmission EM were viewed under a FEI Tecnai Spirit microscope (Arizona Health Science Center Core Imaging Facility) operated at 100 kV. Eight-bit TIFF images were captured using an AMT 4 Mpixel camera. Sections for scanning EM were gold coated and viewed under a FEI Company Inspect S microscope operated at 30 kV.

### Cholesterol crystal imaging

Free floating brain sections were washed in 0.1M PBS 3 times for 5 minutes per wash. Sections were mounted and dried on a slide warmer at 40°C for 1 hour. Sections were then washed 3 times in distilled water for 2 minutes and dried on a slide warmer. Sections were then coverslipped using glycerol, and the edges of the coverslips were sealed with clear nail polish. Cholesterol crystals in the infarct were visualized with standard polarized light microscopy using a Keyence BZ-X700 microscope. Images of the infarct that contained cholesterol crystals were captured using a 20x objective. Lipofuscin autofluorescence was captured as an overlay using a Texas Red filter cube to localize the lesion.

### Cholesterol crystal isolation and verification

Sucrose density centrifugation was performed to verify that the crystals visualized in the area of liquefaction following stroke have the correct density for cholesterol crystals. Stroked C57BL/6 mice (n=5-10) were euthanized 7 weeks following DH stroke, as described above. The infarcts and contralateral cortices were dissected and each dissected region was pooled. The tissue samples were then placed into two sterile 15 mL conical centrifuge tubes containing 750 µL of media (serum-free Neurobasal media, 1× B27 supplement, L-glutamine, and penicillin/streptomycin, without protease/phosphatase inhibitors; Thermo Fisher Scientific). Pooled tissue was then gently triturated by passage through a sterile P1000 pipet. Dissociated cells were centrifuged at 300 rpm for 10 minutes at 4°C. The supernatant was removed, the pellet was re-suspended in 750 µL of fresh media followed by gentle trituration with a P1000 pipet, and the sample re-centrifuged at 300 rpm for 10 minutes at 4°C. Again, the supernatant was removed, the pellet was re-suspended in fresh media, and stored at −80°C. Prior to pouring the sucrose density gradient, infarct samples were spun at 4,000 rpm for 15 minutes at 4°C. The supernatant was extracted and placed in a 2 mL screw cap micro tube. 1 mL of lysis buffer containing protease and phosphatase inhibitors was added to the pellet and the sample was sonicated in bursts. Following sonication, samples were spun at 13,000 rpm for 15 minutes at 4°C. The supernatant was extracted and placed in a 2 mL screw cap micro tube and the pellet was resuspended in 200 µL of lysis buffer and triturated. Each sample was poured into a linear sucrose gradient (0-2 M sucrose). Gradients were then centrifuged at 38,000 x g for 3 hours in an ultracentrifuge. Fractions were removed in 200 µL aliquots from the top of the gradient and analyzed for sucrose content with a refractometer. Using polarized microscopy, 10 µL of each fraction was pipetted onto a slide and the number of cholesterol crystals present in each fraction was counted. This crystal isolation experiment was repeated in three separate experiments. To determine if the formation of cholesterol crystals is specific to the brain following ischemic injury, the above procedure was repeated on heart tissue from mice euthanized at 4 and 8 weeks following MI.

### Quantification of neuronal degeneration

Neurodegeneration in OPN^-/-^ mice was measured by quantifying NeuN in the peri-infarct area. We used NIH Image J analysis software to perform global thresholding in conjunction with a watershed algorithm on both the ipsilateral and equivalent location on the contralateral hemisphere. Fields covering the peri-infarct area were analyzed at bregma 0.02 mm using a 40X objective. Data is presented as the percent area occupied by NeuN immunoreactivity in the ipsilateral/contralateral hemisphere.

### Ladder rung test

Ladder rung testing was performed on WT and OPN^-/-^ mice to assess changes in motor recovery following stroke, as previously described (Pollak et al., 2012). The test was performed at baseline, twice a week for the first 4 weeks post-surgery, and once a week between 5 and 7 weeks post-surgery. A plexiglass ladder measuring 30 inches across with rungs 0.25 inches apart, was supported on side platforms. Traversing animals were recorded from below the ladder with a handheld video recorder. Mice underwent four training sessions (two traverses across the ladder per session) on the ladder before testing commenced. For each testing session, mice traversed the ladder twice. Each video was scored for correct and missed steps. Data is presented as percent correct foot placement versus total foot placement.

### Statistical analysis

Multiplex immunoassays and comparisons between mutant mice and their WT controls were performed with blinding to experimental condition. However, blinding was not possible for immunostaining experiments because of the infarcts being visible in stroked and MI mice. Statistical analyses were performed with Prism 6.0 software (GraphPad). Data are expressed as mean ± standard error of the mean (SEM). Group sizes and statistical tests for each experiment are provided in Table 1.

**Table 1:**
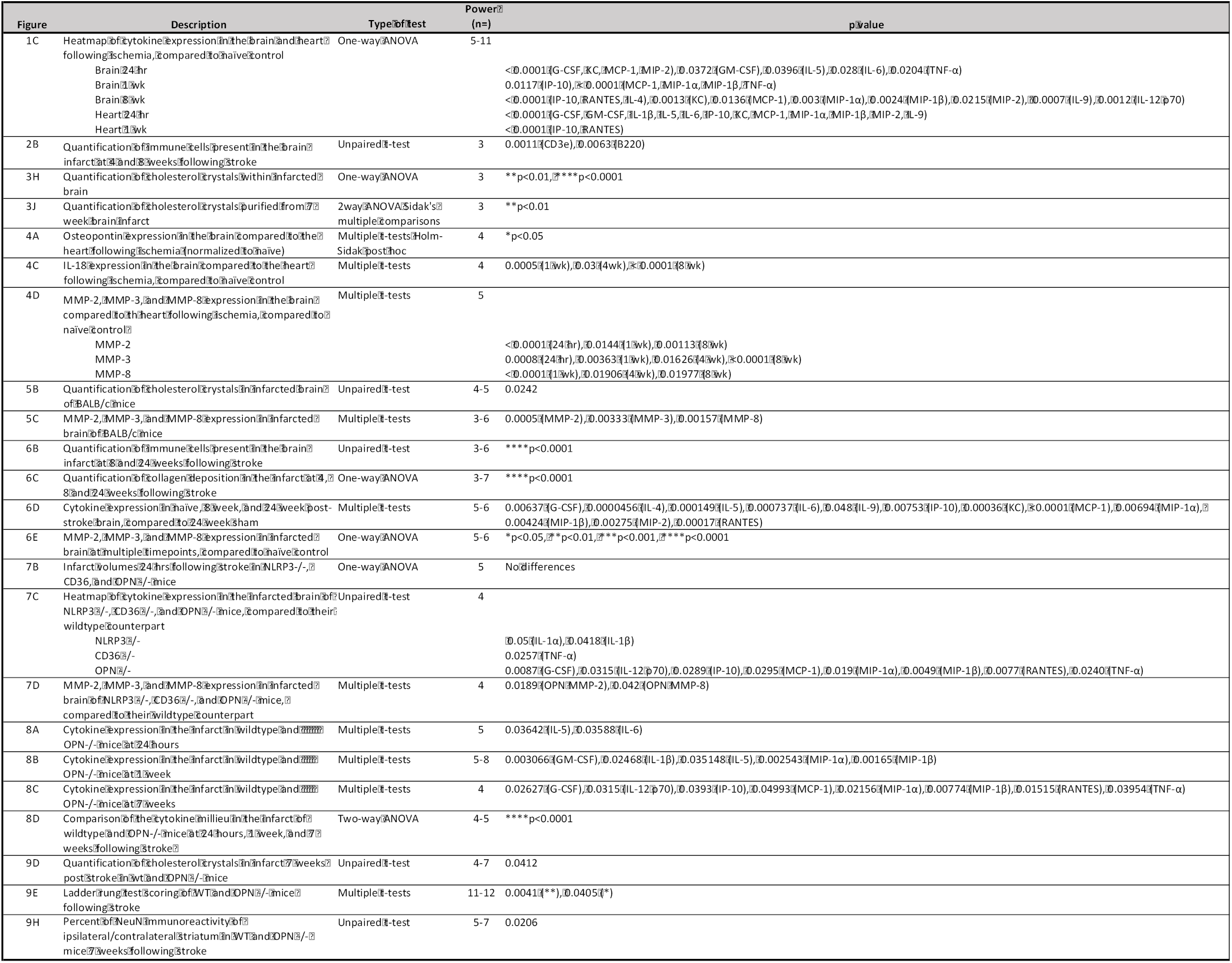
Statistical table

## Results

We have previously demonstrated that liquefactive necrosis is imperfectly segregated from uninfarcted brain tissue by glial scars and have provided evidence that it intensifies post-stroke injury in the weeks after stroke (Zbesko et al., 2018). However, the reason why the brain undergoes liquefactive necrosis in response to ischemia, and why the infarct remains in a liquefactive state for weeks following stroke is unknown. Therefore, in order to improve our understanding of why the brain responds to ischemia by liquefactive necrosis rather than coagulative necrosis, which is the more typical response to ischemia that occurs in other tissues (Robbins et al., 2010), we first compared the kinetics and characteristics of the inflammatory response to ischemia in both the brain and the heart. Mice underwent experimentally induced stroke or MI and were sacrificed 24 hours, 1 week, 4 weeks, and 8 weeks later (**Figure 1A**). Tissue was processed for multiplex immunoassay, histology, and immunostaining.

**Figure 1.**
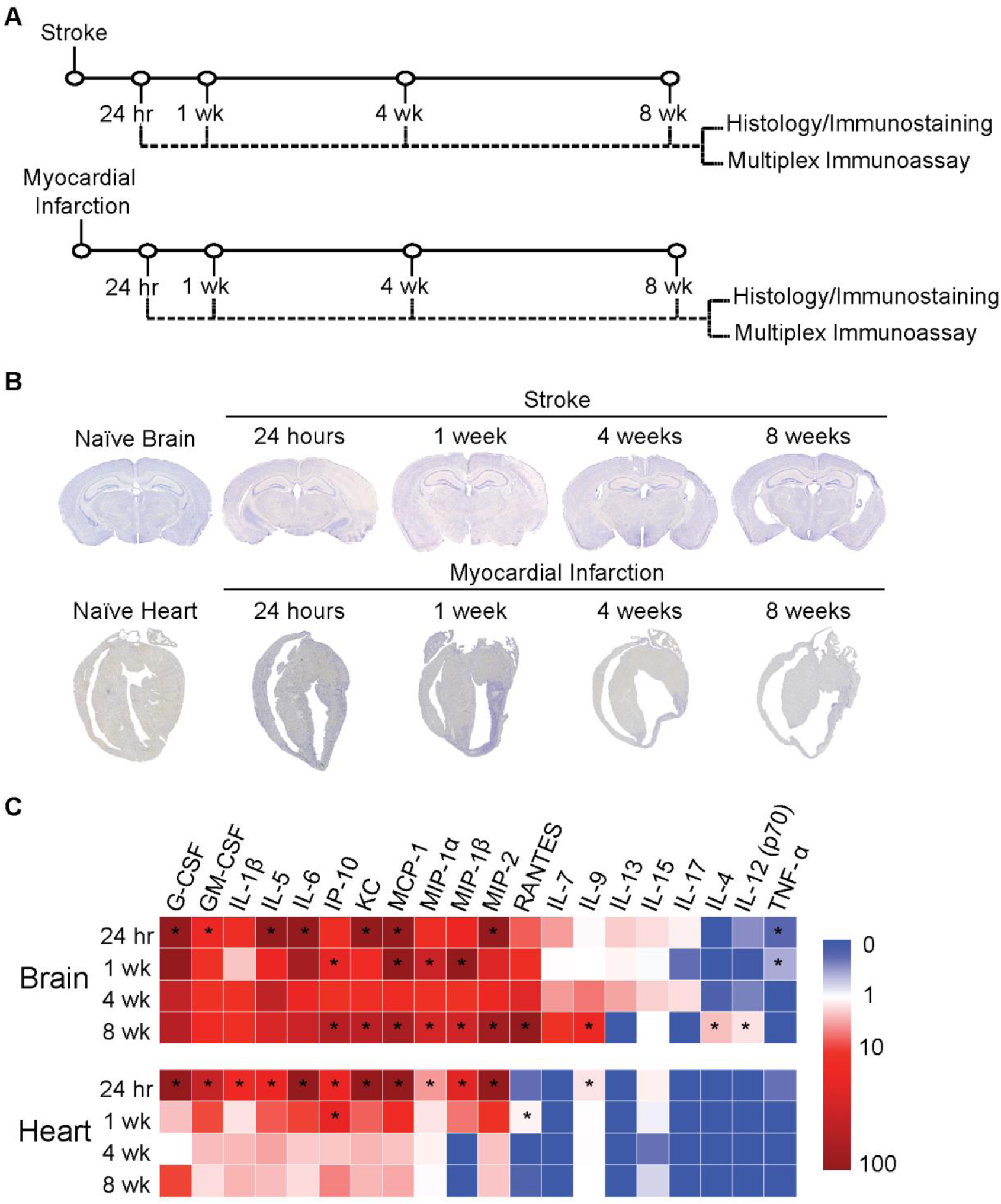
The inflammatory response to ischemic injury is slower to resolve in the brain compared to the heart. (**A**) Experimental design. Mice were either given a stroke or myocardial infarction and were euthanized 24 hours (hr), 1 week (wk), 4 weeks, or 8 weeks following surgery. Infarcted tissue was dissected and processed for analysis by either histology and immunostaining or multiplex immunoassay. (**B**) Nissl staining demonstrates tissue morphology following the respective models of ischemia in the brain and heart. (**C**) Heat map of inflammatory marker expression detected by multiplex immunoassay in the brain and heart at 24 hours, 1 week, 4 weeks, and 8 weeks following ischemia. Values are expressed as fold change relative to naïve heart and brain controls. Group sizes and statistical tests are provided in Table 1.

Each model of ischemia produced a large infarct of comparable size, and as expected, there was extensive tissue remodeling in both organs in the weeks following ischemia (**Figure 1B**). However, although the acute cytokine responses were broadly similar within the infarcted tissue in both the brain and the heart, the chronic inflammatory responses differed markedly. In both tissues, the initial cytokine response subsided within 4 weeks. However, a second wave of pro-inflammatory cytokine expression appeared between weeks 4 and 8 in the brain. This was evidenced by a significant increase in the levels of multiple inflammatory cytokines including IP-10, KC, MCP-1, MIP-1α, MIP-1β, MIP-2, RANTES, IL-9, IL-4, and IL-12 (p70) within the infarct at 8 weeks post stroke when compared to naïve brain tissue (**Figure 1C**).

Furthermore, although there was substantial immune cell infiltration in both the brain and heart at 1 week following ischemia (data not shown), at 4 and 8 weeks following ischemia, there was a clear discrepancy in the amount of immune cell infiltration and fibrosis present within each type of infarct that was easily discernible by visual inspection (**Figure 2**). There was also a significant increase in both CD3e (T-lymphocyte) and B220 (B-lymphocyte) immunoreactivity between 4 and 8 weeks following ischemia in the brain (**Figure 2A-B**), with no apparent increase in collagen deposition in the brain infarcts between 4 weeks and 8 weeks following ischemia (**Figure 2A**). In comparison to the brain, the heart infarcts contained markedly more collagen (**Figure 2C**) and substantially fewer T-lymphocytes, B-lymphocytes, and CD68+ macrophages at 4 weeks and 8 weeks following ischemia (**Figure 2C-D**). These data indicate that liquefaction of the brain following stroke correlates with a second wave of cytokine expression, reduced collagen deposition, and more extensive chronic immune cell infiltration than occurs in the heart following a MI.

**Figure 2.**
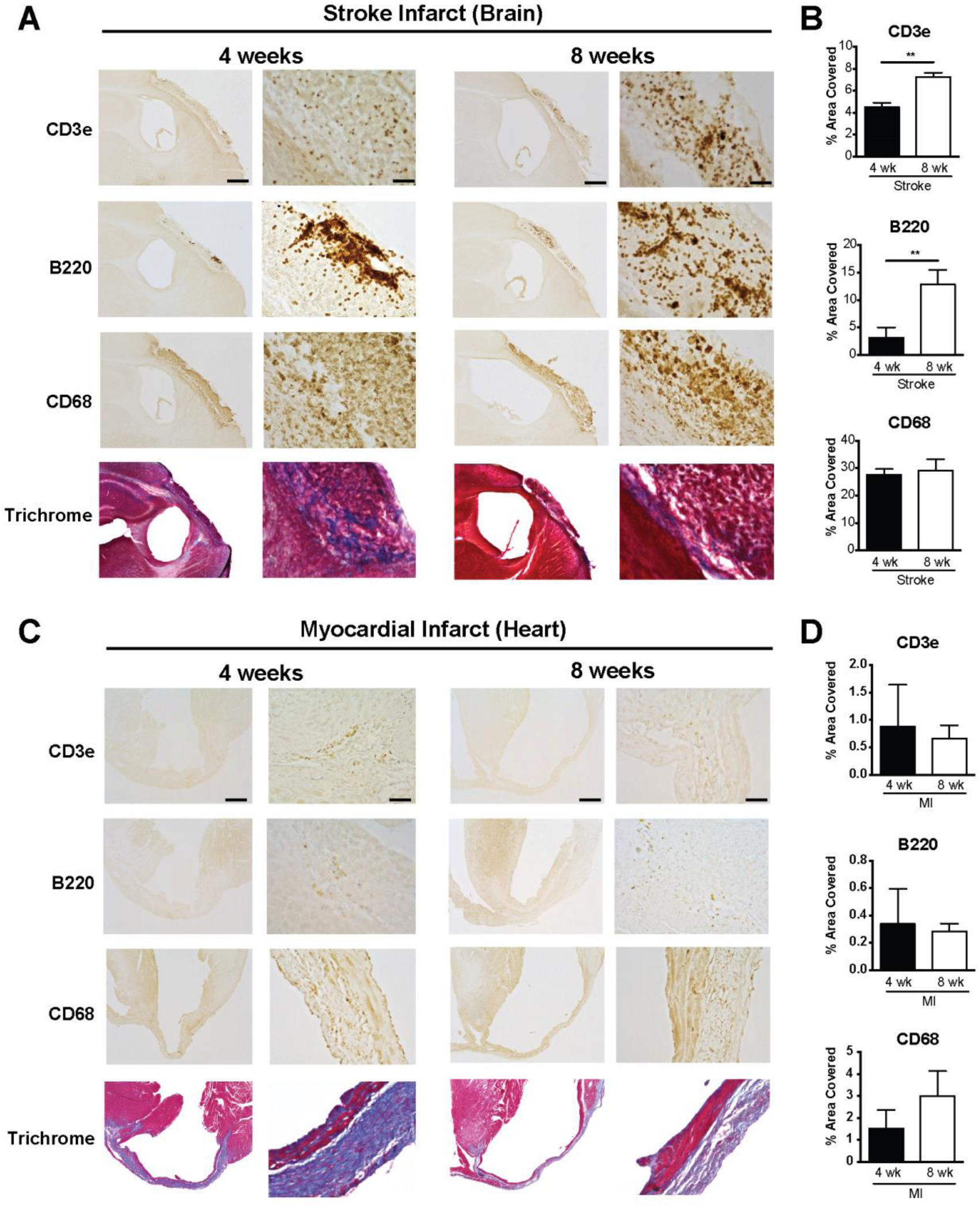
Chronic immune cell infiltration is more extensive in response to ischemic injury in the brain compared to the heart. (**A**) Representative low and high magnification images of T-lymphocytes (CD3e), B-lymphocytes (B220), microglia/macrophages (CD68), and collagen (blue in the trichrome stain) in the area of liquefactive necrosis in the brain at 4 and 8 weeks following ischemic injury. Scale bar, 500 µm for low (4x) magnification images, and 50 µm for high (20x) magnification images. (**B**) Percent area covered by the different immune cell markers in the area of liquefactive necrosis in the brain at 4 and 8 weeks following ischemic injury. Data represented as mean ± SEM. Group sizes and statistical tests are provided in Table 1. (**C**) Representative low and high magnification images of T-lymphocytes (CD3e), B-lymphocytes (B220), microglia/macrophages (CD68), and collagen (blue in the trichrome stain) in the infarct in the heart at 4 and 8 weeks following ischemic injury. Scale bar, 500 µm for low (4x) magnification images, and 50 µm for high (20x) magnification images. (**D**) Percent area covered by the different immune cell markers in myocardial infarcts at 4 and 8 weeks following ischemic injury. Data represented as mean ± SEM. Group sizes and statistical tests are provided in Table 1. There is more extensive infiltration of T-lymphocytes, B-lymphocytes, and microglia/macrophages in the brain compared to the heart. Conversely, there is much more collagen deposition in the heart compared to the brain.

To understand why the second wave of cytokine expression develops in the infarct in the weeks following stroke, and why the chronic immune cell infiltrate is more substantial and sustained, we looked closely at the configuration of cytokines that were significantly elevated at the 8-week time point (**Figure 1C**). This revealed a molecular fingerprint reminiscent of the cytokine profile present in atheroma. Specifically, IP-10, KC, MCP-1, MIP-1α, MIP-1β, MIP-2, RANTES, IL-9, IL-4, and IL-12 (p70) have all been found in atherosclerotic plaques (Braunersreuther et al., 2007; Ramji and Davies, 2015). Significantly, blocking IP-10 and MCP-1 results in atherosclerosis regression in mice (Braunersreuther et al., 2007). Our previous data demonstrate that the human brain at the stage of liquefactive necrosis also contains increased levels of IP-10, MCP-1, MIP-1α, MIP-1β, and IL-12 (p70) (Nguyen et al., 2016). Therefore, liquefaction of the brain following stroke shares several cytokine characteristics with atherosclerosis.

In light of this overlap in the cytokine profile between liquefactive necrosis of the brain following stroke and atherosclerosis, we hypothesized that the sustained state of liquefactive necrosis in response to stroke may be caused by excessive cholesterol within phagocytic cells that engulf cholesterol rich myelin debris. Therefore, to first provide evidence that there is a pool of myelin debris in the area of the infarct that undergoes liquefactive necrosis, we performed Fluoro-Jade C staining on tissue at 24 hours, 1 week, 4 weeks, and 8 weeks following stroke. As expected, this demonstrated that there is extensive neurodegeneration in the infarcted tissue 24 hours following stroke, which progresses for at least 1 week following stroke (**Figure 3A**).

**Figure 3.**
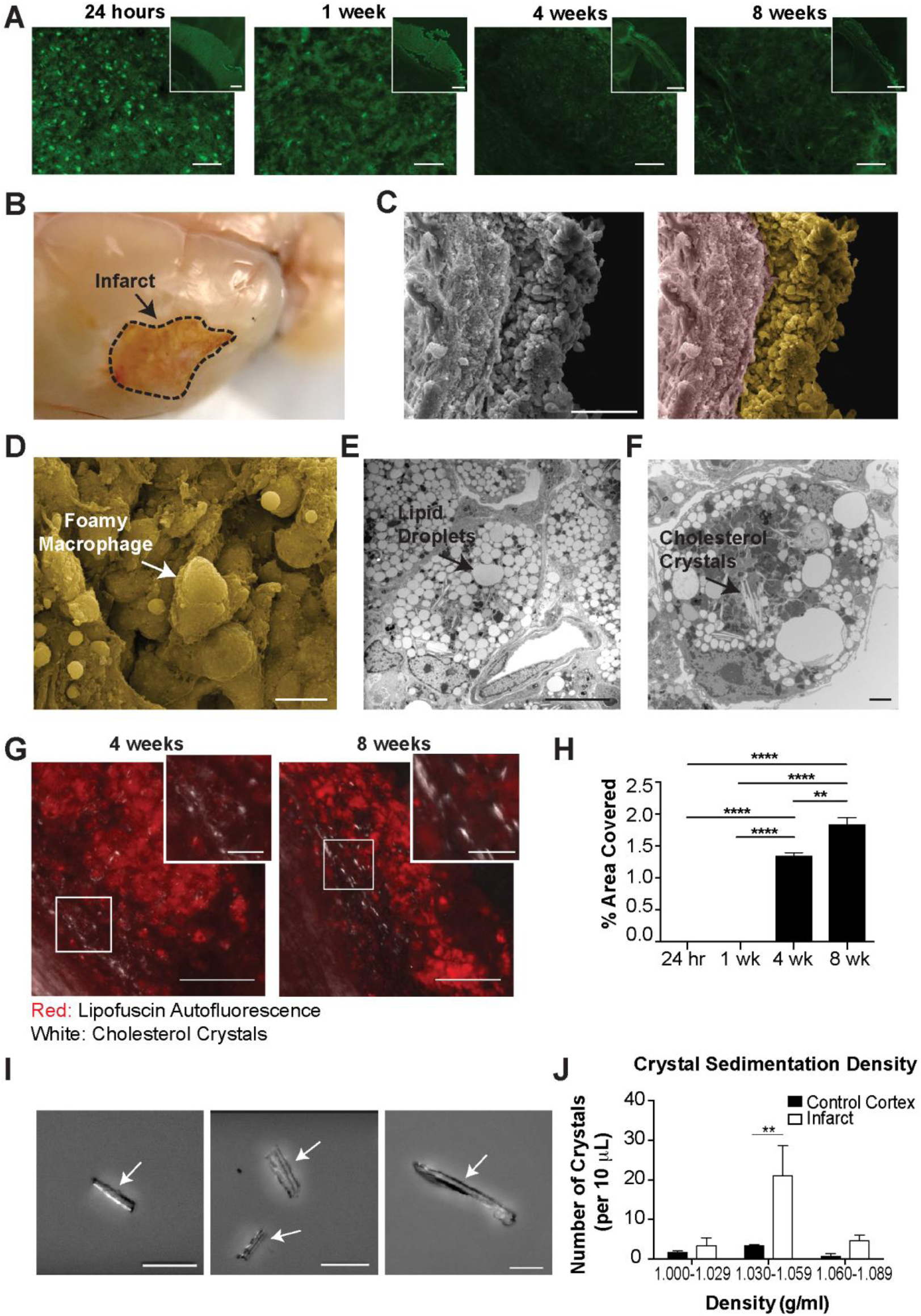
Foamy macrophages within areas of liquefactive necrosis contain cholesterol crystals. (**A**) Representative images of Fluoro-Jade C staining show degenerating neurons in the lesion 24 hours, 1 week, 4 weeks, and 8 weeks following stroke. Scale bar 50 µm. Inset scale bar 500 µm. (**B**) Gross pathology of a lesion (delineated) 7 weeks following stroke. The infarcted area is in a liquefactive state (arrow). (**C**) Original (left image) scanning EM image of a 7 week infarct with a dense population of cells (pseudocolored yellow; right image) and glial scar (pseudocolored purple; right image). Scale bar, 100 µm. The infarct is filled with foamy macrophages (arrow), as seen magnified in (**D**). Scale bar, 50 µm. (**E**) Transmission EM of a macrophage in the infarct at 7 weeks following stroke. There is an accumulation of lipid droplets (arrow), providing the macrophage with its characteristic foamy appearance. Scale bar, 10 µm. (**F**) Transmission EM of a foamy macrophage in the infarct at 7 weeks following stroke with putative cholesterol crystal clefts in the cytoplasm (arrow). Scale bar, 2 µm. (**G**) Confirmation of the presence of cholesterol crystals (white) in the infarct by overlaying images taken with polarized and fluorescence microscopy. Lipofuscin autofluorescence (red) demarcates the area of infarction. Scale bar, 100 µm. Inset scale bars, 25 µm. (**H**) Quantification of cholesterol crystals in the lesion at 24 hours, 1 week, 4 weeks, and 8 weeks following stroke. Data represents mean ± SEM. (**I**) Cholesterol crystals, purified from 7 week infarcted brains using sucrose density gradient centrifugation, and visualized under polarized microscopy. Cholesterol crystals (denoted by arrows) were identified by the presence of light refraction and display a distinct needle and plate-like morphology. Scale bar, 50 µm. (**J**) The number of cholesterol crystals present in fractionated densities from chronic stroke infarcts or the equivalent area on the contralateral hemisphere (control cortex). Data represent mean ± SEM from n=3 sucrose density gradient centrifugation runs with n=5-10 mice per experiment. Group sizes and statistical tests are provided in Table 1.

We next evaluated the macroscopic (**Figure 3B**) and microscopic (**Figure 3C-D**) morphological features of liquefactive necrosis and looked for the presence of cholesterol crystals, which are a hallmark of excessive cholesterol in a biological system (**Figure 3E-J**). As expected, scanning EM revealed that liquefactive necrosis is predominantly comprised of a sea of foamy macrophages morphologically similar to the foam cells found in atherosclerotic lesions (**Figure 3C-D**). Transmission EM revealed that each macrophage contains an abundance of lipid droplets (**Figure 3E**). Importantly, transmission EM also provided evidence that many of these macrophages contain cholesterol crystals (**Figure 3F**).

Cholesterol crystals are birefringent and change the polarization of transmitted light (Kashiwagi et al., 2014). Therefore, to validate that macrophages in areas of liquefactive necrosis contain cholesterol crystals, we imaged the 24 hours, 1 week, 4 week, and 8 week-old stroke infarcts using polarized light microscopy. This revealed that birefringent crystals appear within stroke infarcts between weeks 1 and 4 following stroke and are even more abundant at 8 weeks following stroke (**Figure 3G-H**). This timeline of appearance of crystals correlates with the second wave of cytokine expression within the infarct and the failure of the immune cell infiltrate to resolve between 4 and 8 weeks following stroke, as occurs in the heart.

Although transmission EM and polarized light microscopy are gold standard techniques for the identification of cholesterol crystals (Kashiwagi et al., 2014), we also biochemically identified these crystals as comprised of cholesterol based on their equilibrium density sedimentation. Cholesterol crystals are known to have a density of 1.04 g/ml (Tangirala et al., 1994). Crystals isolated from cells extracted from areas of liquefactive necrosis 7 weeks following stroke had heterogeneous needle-like and plate-like morphologies, which matches the physical description of cholesterol crystals (Tangirala et al., 1994) (**Figure 3I**), and they sedimented in fractions ranging from 1.030-1.059g/ml, in agreement with the expected density of cholesterol crystals (**Figure 3J**). Density centrifugation of heart tissue at 8 weeks following MI failed to isolate any birefringent crystals with the reported density of cholesterol crystals. These experiments provide evidence that chronic liquefactive necrosis of the brain in response to stroke is associated with the accumulation of cholesterol crystals within macrophages, and that cholesterol crystals do not accumulate in the heart following MI.

To determine if liquefaction of the brain following stroke shares other characteristics with atherosclerosis we measured the levels of OPN, IL-18, MMP-2, MMP-3, and MMP-8. The rationale for measuring OPN is that it is one of the most abundant proteins present within atheroma, and is expressed at high levels in other crystallopathies (Khan et al., 2002). The rationale for measuring IL-18 is that cholesterol crystals activate the NLRP3 inflammasome in atheroma, and IL-18 is a hallmark of inflammasome activation (Duewell et al., 2010; Baldrighi et al., 2017; Hoseini et al., 2017; Karasawa and Takahashi, 2017). The rationale for evaluating levels of MMP-2, MMP-3, and MMP-8 is that cholesterol crystals induce MMP expression in macrophages, and MMPs are abundant in atheroma (Beaudeux et al., 2004; Watanabe and Ikeda, 2004; Lenglet et al., 2013; Corr et al., 2016). Multiplex immunoassay analyses revealed that OPN is present in areas of liquefactive necrosis for at least 8 weeks following stroke (**Figure 4A**). Immunofluorescence confirmed that it colocalizes with IBA1+ microglia/macrophages as previously reported (Gliem et al., 2015) (**Figure 4B**). IL-18 is also present in the area of liquefactive necrosis for at least 8 weeks following stroke (**Figure 4C**), and there are also strikingly high levels of MMP-2, MMP-3, and MMP-8 (**Figure 4D**). By contrast, OPN, IL-18, MMP-2, MMP-3, and MMP-8 were present at substantially lower levels at chronic time points following MI (**Figure 4A, C-D**).

**Figure 4.**
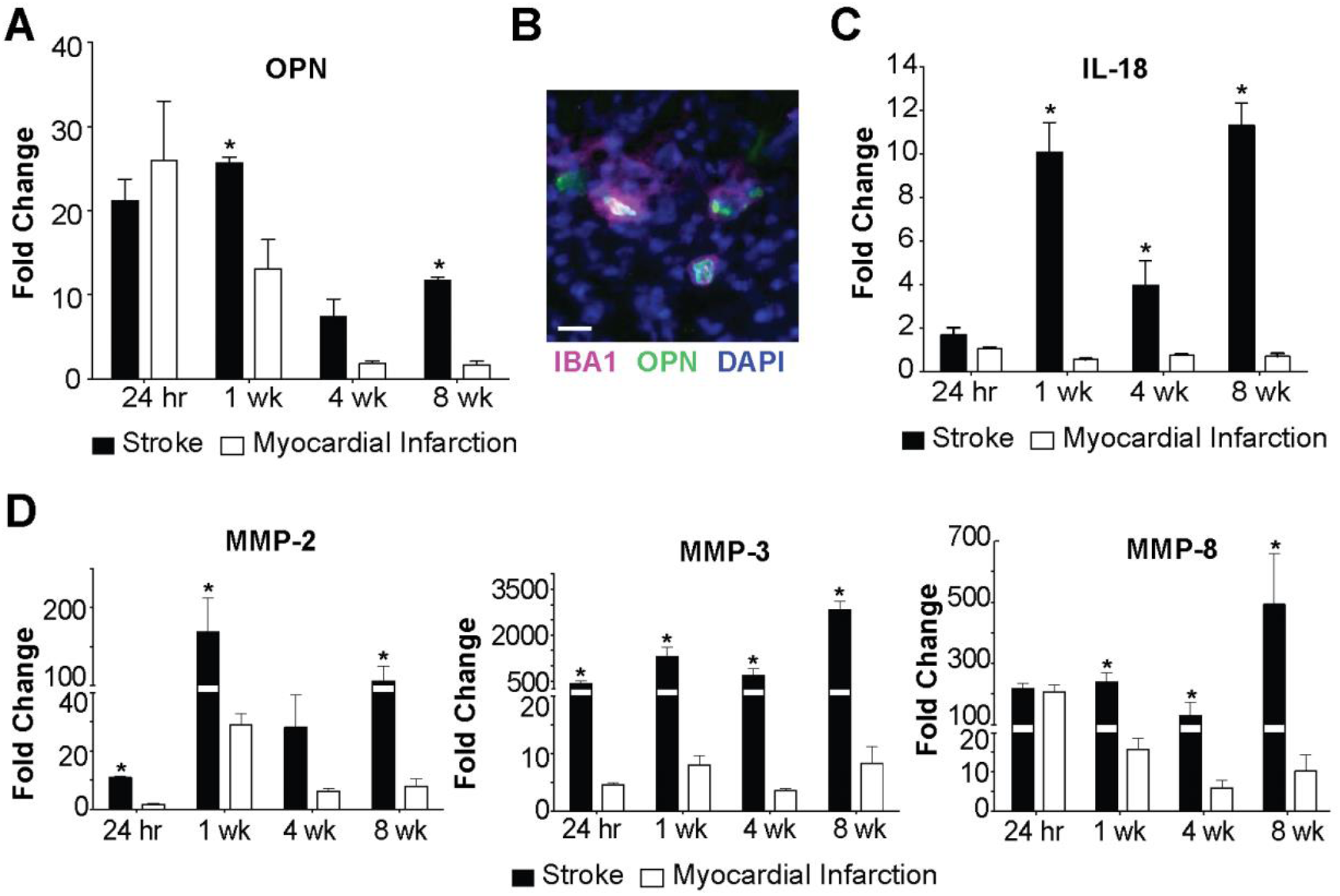
Liquefaction of the brain following stroke shares multiple characteristics with atherosclerosis. (**A**) OPN levels are significantly elevated in brain infarcts compared to heart infarcts at 1 week and 8 weeks following ischemic injury. Data represent fold changes relative to naïve brain or heart tissue ± SEM. *p<0.05 compared to heart. (**B**) OPN colocalizes with IBA1+ cells in the brain infarcts at 7 weeks following stroke. Image representative of 5 mice. Scale bar, 20 µm. (**C**) IL-18 levels are significantly increased in brain infarcts compared to heart infarcts at 1 week, 4 weeks, and 8 weeks following ischemic injury. Data represent fold changes relative to naïve brain or heart tissue ± SEM. (**D**) MMP-2 levels were significantly elevated within the brain infarcts at 24 hours, 1 week, and 8 weeks following ischemic injury compared to the heart infarcts. MMP-3 levels were significantly elevated within the brain infarct at 24 hours, 1 week, 4 weeks, and 8 weeks following ischemic injury compared to the heart infarcts. MMP-8 levels were significantly elevated within the brain infarcts at 1 week, 4 weeks, and 8 weeks following ischemic injury compared to the heart infarcts. Data represent fold changes relative to naïve brain or heart tissue ± SEM. Group sizes and statistical tests are provided in Table 1.

We next tested if the formation of cholesterol crystals and the high expression of MMPs in areas of liquefactive necrosis is specific to C57BL/6 mice or specific to DH stroke. BALB/c mice underwent DMCAO without hypoxia and were sacrificed 7 weeks later. These mice also developed birefringent crystals in the infarct in the weeks after stroke (**Figure 5A-B**), and had abundant expression of MMP-2, MMP-3, and MMP-8 (**Figure 5C**) in the area of liquefaction 7 weeks following stroke. The formation of cholesterol crystals in chronic stroke infarcts is not mouse strain or stroke model dependent.

**Figure 5.**
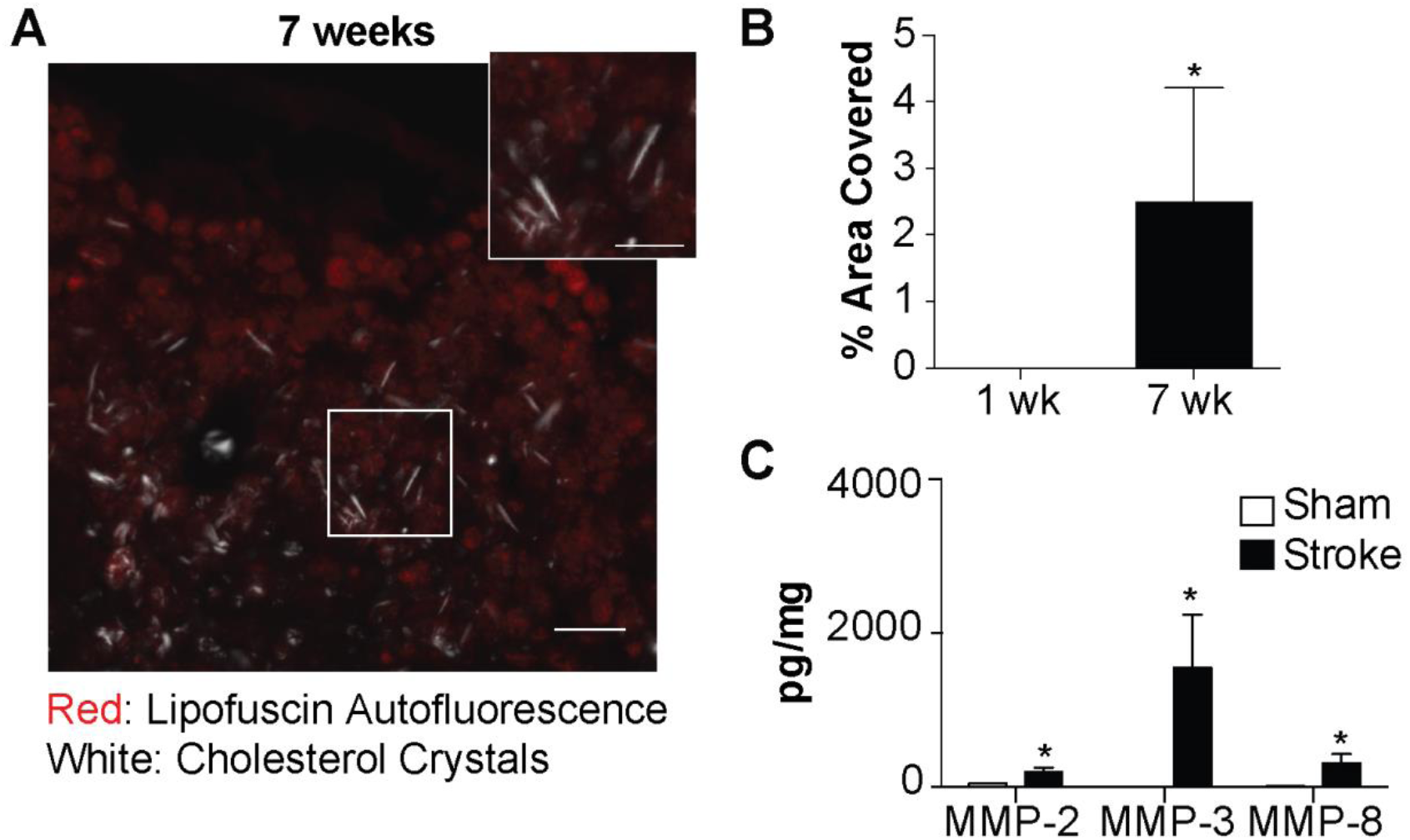
Cholesterol crystals and MMPs also accumulate in the area of liquefactive necrosis following stroke in BALB/c mice. (**A**) Representative image of cholesterol crystals present in the infarct of BALB/c mice 7 weeks following stroke, visualized using polarized microscopy overlaid with fluorescence microscopy. Lipofuscin autofluorescence (red) delineates the lesion. Scale bar, 50 µm. Inset scale bar, 25 µm. (**B**) Cholesterol crystals accumulate in the infarct of BALB/c mice between 1 week and 7 weeks following stroke. (**C**) Levels of MMP-2, MMP-3, and MMP-8 are significantly increased in the infarct of stroked BALB/c mice compared to an equivalent area of the cortex in sham mice. Group sizes and statistical tests are provided in Table 1.

Even though liquefactive necrosis is imperfectly contained by glial scarring and may intensify post-stroke injury (Zbesko et al., 2018), it is currently unknown how long it takes for liquefactive necrosis to resolve in the brain after stroke. Therefore, C57BL/6 mice underwent DH stroke and were sacrificed 24 weeks later. Immunostaining revealed a substantial reduction in CD3e+ T-lymphocyte, B220+ B-lymphocyte, and CD68+ macrophage/microglia infiltration at 24 weeks following stroke compared to 8 weeks following stroke, and trichrome staining demonstrated a concurrent increase in collagen scarring (**Figure 6A-C**). Multiplex immunoassay determined that cytokine levels in the infarct at 24 weeks were substantially and significantly lower compared to the corresponding area of infarct from mice sacrificed 8 weeks following stroke (**Figure 6D**). Levels of MMP-2, MMP-3, and MMP-8 were also substantially reduced and had returned to baseline levels (**Figure 6E**). Based on these data, liquefactive necrosis is resolved by 24 weeks following stroke in young adult C57BL/6 mice.

**Figure 6.**
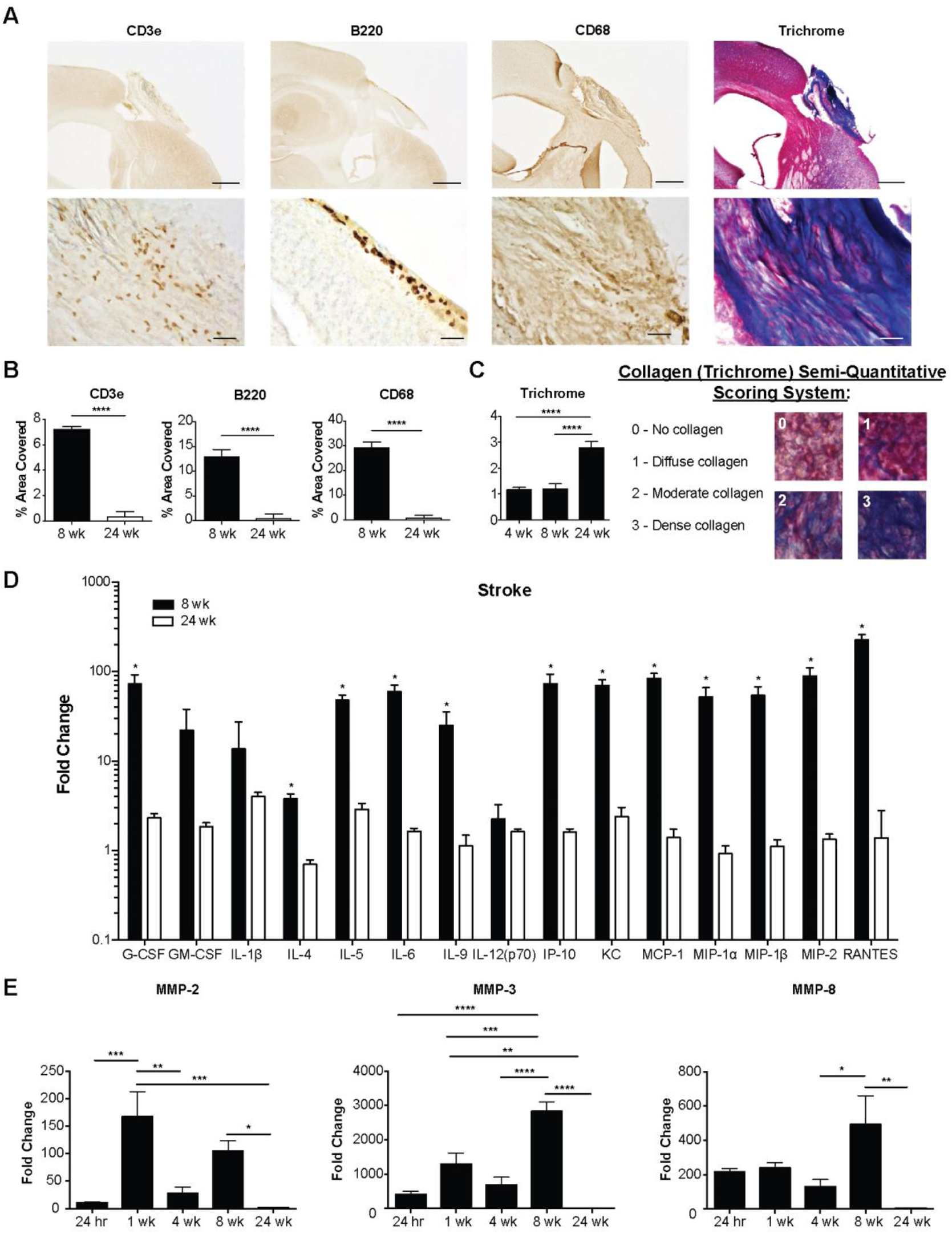
Liquefactive necrosis subsides and a collagen scar is formed by 24 weeks following stroke in C57BL/6 mice. (**A**) Top panels show representative images of immune cell infiltration and collagen deposition in brain infarcts at 24 weeks post-stroke. Scale bar, 500 µm. Bottom panels show higher magnification images. Scale bar, 50 µm. (**B**) Quantification of immune cell infiltration at 8 and 24 weeks post-stroke. There is a decrease in the number of T-lymphocytes (CD3e; left graph), B-lymphocytes (B220; middle graph), and macrophages (CD68; right graph) in the infarct 24 weeks following stroke compared to 8 weeks. (**C**) Quantification of collagen deposition in the infarct at 4, 8 and 24 weeks post-stroke. There is a marked increase in collagen (Trichrome; blue) staining at 24 weeks following stroke compared to 4 and 8 weeks. (**D**) Multiplex immunoassays demonstrate a concomitant decrease in cytokines/chemokines within the infarct at 24 weeks following stroke compared to 8 weeks following stroke. Data presented as fold change from naïve. (**E**) Multiplex immunoassays indicate that MMP levels within the infarct have also returned to baseline levels at 24 weeks following stroke. Data represents mean ± SEM. Group sizes and statistical tests are provided in Table 1.

We then tested if targeting intermediaries in the pathophysiology of atherosclerosis using genetically modified mice could mitigate the production of cytokines and proteases within chronic stroke infarcts and improve stroke outcome. The intermediaries targeted were NLRP3, CD36, and OPN. The rationale for NLRP3 was that we hypothesized that the genetic ablation of NLRP3 would prevent the activation of the NLRP3 inflammasome by cholesterol crystals. CD36 is a scavenger receptor involved in both cholesterol uptake by macrophages and inflammasome activation in atherosclerosis (Stewart et al., 2010; Kagan and Horng, 2013; Park, 2014), and OPN regulates macrophage chemotaxis (Lund et al., 2013), the production of MMPs by macrophages (Mi et al., 2007) and enhancement of phagocytosis via opsonization (Schack et al., 2009) (**Figure 7A**).

**Figure 7.**
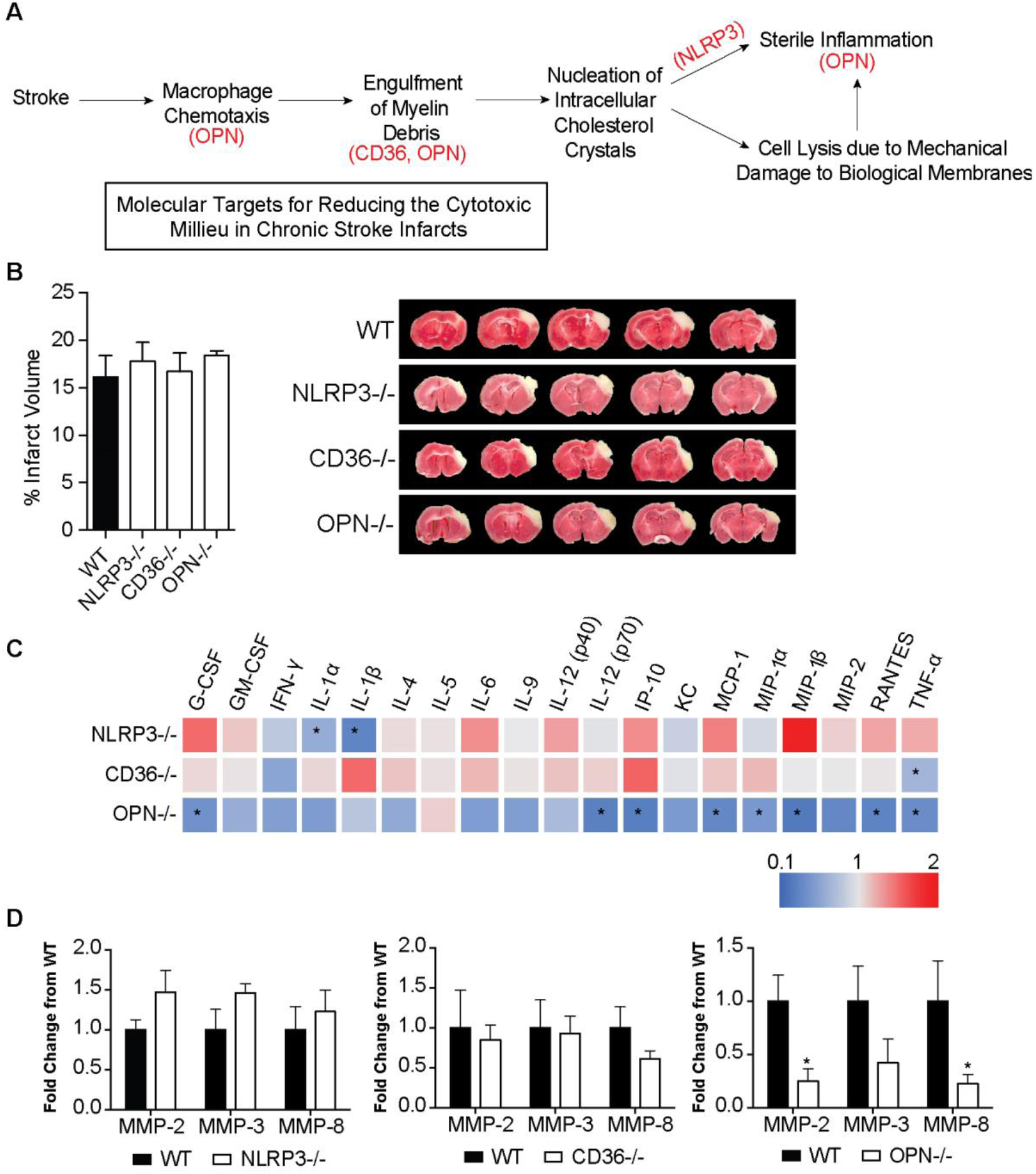
Molecular targets for reducing the cytotoxic milieu in chronic stroke infarcts. (**A**) Rationale for targeting OPN, CD36, and NLRP3 as a means of reducing the inflammatory response in areas of liquefactive necrosis following stroke. (**B**) Infarct volume is equivalent in WT, NLRP3, CD36, and OPN deficient mice at 24 hours following stroke. Data represents mean ± SEM. (**C**) Heat map of the levels of inflammatory markers in infarcted brain obtained by multiplex immunoassays demonstrates that the chronic inflammatory response was attenuated the most in OPN^-/-^ mice at 7 weeks post stroke. NLRP3^-/-^ mice showed a significant reduction in the chronic levels of IL-1α and IL-1β. CD36^-/-^ mice only showed a significant reduction in chronic levels of TNF-α. Data presented as fold change relative to WT mice. (**D**) Multiplex immunoassays of MMPs show no significant changes in MMP levels in the chronic 7 weeks old infarcts of NLRP3^-/-^ (left graph) and CD36^-/-^ (middle graph) mice compared to WT mice. MMP levels are significantly and substantially lower in the chronic infarct of OPN^-/-^ mice (right graph) compared to WT mice. Data represents mean ± SEM. Group sizes and statistical tests are provided in Table 1.

WT, NLRP3^-/-^, CD36^-/-^, and OPN^-/-^ mice underwent DH stroke and were sacrificed 24 hours later for determination of acute infarct size. TTC staining determined that acute infarct volume is equivalent in each strain following the DH stroke model (**Figure 7B**). Congruently, we have previously shown that infarct size following the intraluminal filament MCAO model of stroke is equivalent in OPN^-/-^ and WT mice (Meller et al., 2005), however, NLRP3^-/-^ and CD36^-/-^ mice have been reported to have smaller infarct volumes compared to WT mice following intraluminal filament MCAO (Cho et al., 2005; Kunz et al., 2008; Yang et al., 2014; Garcia-Bonilla et al., 2015). We ascribe this discrepancy in infarct size between models to the DH stroke model having a smaller salvageable penumbra at acute timepoints than the intraluminal filament MCAO stroke model.

Mice then underwent DH stroke and were sacrificed 7 weeks later to evaluate the impact of each gene on the chronic inflammatory response to stroke. The levels of cytokines and MMPs in the area of liquefaction were analyzed by multiplex immunoassay and compared to WT mice. Levels of IL-1α and IL-1β were significantly reduced in the NLRP3^-/-^ mice (**Figure 7C**), however other cytokine levels and MMP levels were unchanged (**Figure 7C-D**). There were no changes in cytokine or MMP levels in the CD36^-/-^ mice except for a reduction in TNF-α (**Figure 7C-D**). The biggest change in the levels of cytokines and MMP levels was evident in the OPN^-/-^ mice. G-CSF, IL-12(p70), IP-10, MCP-1, MIP-1α, MIP-1β, RANTES, TNF-α, MMP-2, and MMP-8 were all significantly and substantially reduced (**Figure 7C-D**).

To follow up on this finding, additional OPN^-/-^ mice underwent DH stroke and were sacrificed 24 hours and 1 week later to evaluate the impact of OPN deficiency on the inflammatory response to stroke at an acute and subacute time point. At 24 hours following stroke, IL-5 and IL-6 were increased in the infarct of the OPN^-/-^ mice compared to the WT mice (**Figure 8A**). At 1 week following stroke GM-CSF, IL-1β, and IL-5 were reduced, and MIP-1α, and MIP-1β were increased in the infarct of the OPN^-/-^ mice (**Figure 8B**). However, as previously shown (**Figure 8C**) at 7 weeks following stroke, OPN deficiency resulted in a much more substantial alteration of the inflammatory response to stroke (**Figure 8C**), with an overall reduction in the cytokine milieu as determined by two-way ANOVA (**Figure 8D**).

**Figure 8.**
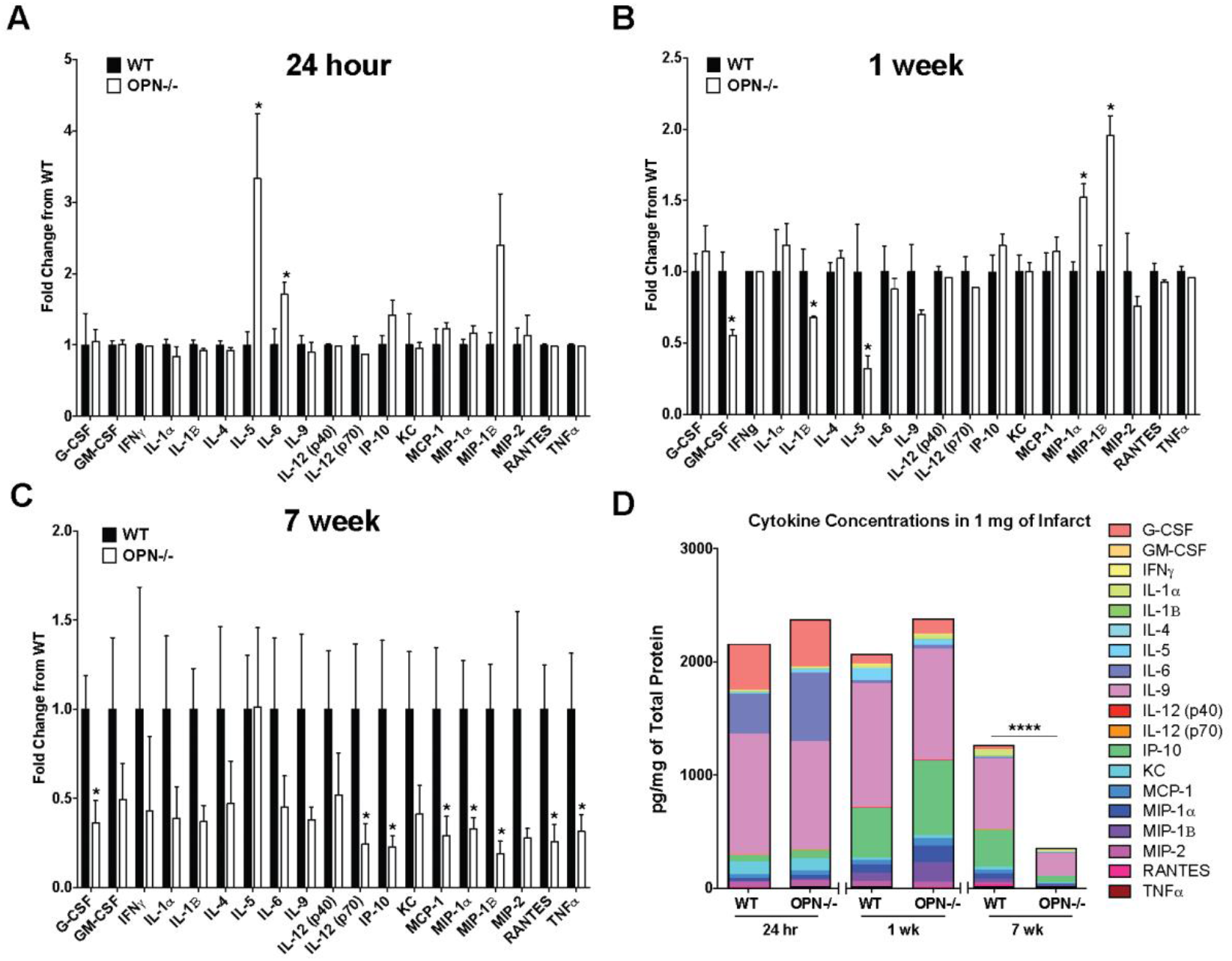
Comparison of the effect of OPN deficiency on the acute, subacute and chronic inflammatory response to stroke. (**A**) At 24 hours following stroke, IL-5 and IL-6 are increased in the infarct of OPN^-/-^ mice compared to WT mice. Data represents mean ± SEM. (**B**) At 1 week following stroke GM-CSF, IL-1β, and IL-5 are reduced, and MIP-1α, and MIP-1β are increased in OPN deficient mice. Data represents mean ± SEM. (**C**) At 7 weeks following stroke, OPN deficiency results in a significant reduction of G-CSF, IL-12(p70), IP-10, MCP-1, MIP-1α, MIP-1β, RANTES, and TNFα. Data represents mean ± SEM. (**D**) The global cytokine response is significantly reduced in OPN^-/-^ at 7 weeks, but not 24 hours or 1 week following stroke by two-way ANOVA. All data represent mean ± SEM. Group sizes and statistical tests are provided in Table 1.

Based on the considerable reduction in cytokine and protease expression present in the area of liquefaction in the OPN^-/-^ mice at 7 weeks following stroke, we next determined if this correlated with an alteration in immune cell infiltration, astrogliosis, and cholesterol crystal accumulation. The number of T-lymphocytes and B-lymphocytes in the infarct of the WT and OPN^-/-^ mice was not significantly different at 7 weeks following stroke (**Figure 9A-B**). Furthermore, there were no significant differences in the amount of GFAP expressed in the peri-infarct location (**Figure 9A-B**). However, cholesterol crystal accumulation was increased in the infarct of the OPN^-/-^ mice compared to the WT mice (**Figure 9C-D**).

**Figure 9:**
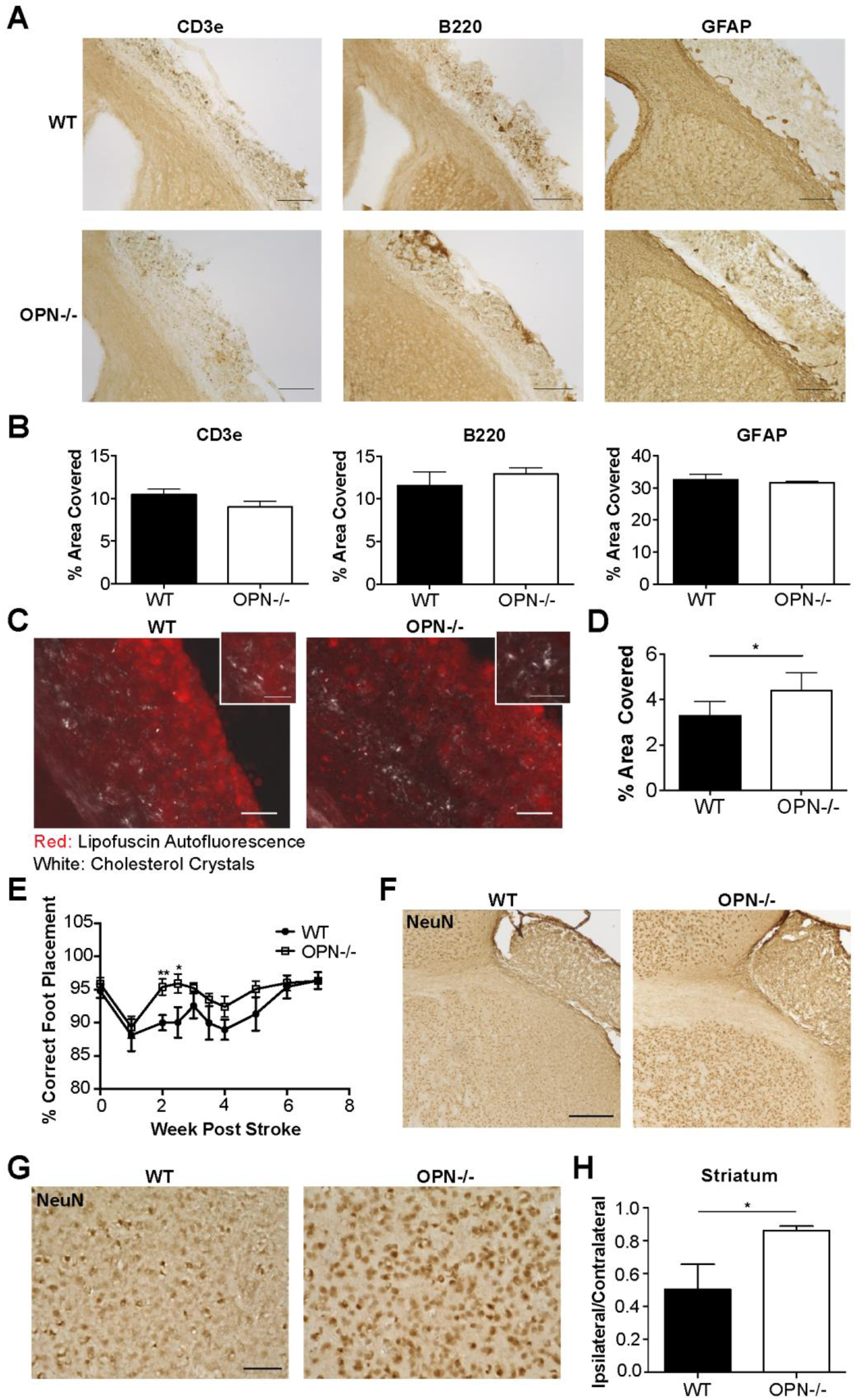
Genetic ablation of OPN improves recovery and reduces secondary neurodegeneration following stroke. (**A**) Representative images of T-lymphocytes (CD3e; left panel), B-lymphocytes (B220; middle panel), and glial scarring (GFAP; right panel) present in the area of liquefactive necrosis in WT (top panel) and OPN^-/-^ (bottom panel) mice 7 weeks following stroke. Scale bar 200 µm. (**B**) Quantification of CD3e, B220, and GFAP immunoreactivity in the infarct of WT and OPN^-/-^ mice 7 weeks following stroke. Data represents mean ± SEM. (**C**) Cholesterol crystals present in the infarct 7 weeks following stroke in WT (left) and OPN^-/-^ (right) mice visualized with polarized microscopy. Lipofuscin autofluorescence (red) delineates infarct. Scale bar 50 µm. Inset scale bar 25 µm. (**D**) Quantification of cholesterol crystals in the infarct of wt and OPN^-/-^ mice 7 weeks following stroke. Data represents mean ± SEM. (**E**) OPN^-/-^ mice were able to recover faster on the ladder rung test compared to WT mice. Ladder rung tests were scored twice per week for the first 4 weeks following stroke and weekly thereafter. (**F**) Representative images showing preservation of neuronal marker NeuN immunoreactivity in the peri-infarct area of OPN^-/-^ mice compared to WT mice at 7 weeks following stroke. Scale bar, 250 µm. (**G**) Higher magnification representative images showing preservation of NeuN immunoreactivity in the peri-infarct area of OPN^-/-^ mice. Scale bar, 50 µm. (**H**) Quantitation of NeuN immunoreactivity in the striatum shows a significant increase in the OPN^-/-^ mice. Data represents mean ± SEM. Group sizes and statistical tests are provided in Table 1.

Finally, to determine how the altered inflammatory response in the area of liquefaction impacts recovery in OPN^-/-^ mice, recovery of motor function was evaluated using a horizontal ladder, and secondary neurodegeneration was evaluated using the neuronal marker, NeuN. OPN^-/-^ mice recovered motor function significantly faster following DH stroke than WT mice (**Figure 9E**) and exhibited a greater preservation of neuron number in the peri-infarct location, as evidenced by increased NeuN immunoreactivity (**Figure 9F-H**).

## Discussion

Using mouse models of MI and ischemic stroke, we tested whether the brain takes longer to heal from ischemic injury than the heart. Using these models, we demonstrated that the inflammatory response to ischemic injury in the brain takes longer to resolve than the inflammatory response to a similar injury in the heart. Further, in the brain we observed a biphasic cytokine response to ischemia, with the second phase initiated between 4 and 8 weeks after ischemic injury. In common with the pathogenesis of atheroma, the second phase included a predominance of foamy macrophages and the presence of T-lymphocytes. The second phase of inflammation also coincided with the accumulation of intracellular and extracellular cholesterol crystals within the infarct.

Cholesterol crystals can activate the NLRP3 inflammasome. Accordingly, we found evidence of NLRP3 inflammasome activity within ischemic stroke infarcts during the chronic stage of liquefactive necrosis by way of a significant elevation of IL-18 at 4 and 8 weeks following stroke. In contrast, elevated IL-18 was not detected in the heart following MI. We also examined MMP expression given the known association of MMPs with atherosclerotic plaque, and their potential participation in liquefaction of the necrotic tissue. There was chronic and abundant expression of MMP-2, MMP-3, and MMP-8 in the area of liquefactive necrosis following stroke compared to the equivalent area of infarction in the heart. Indeed, expression of MMP-2 was 100-fold higher (4,273 pg/mg), expression of MMP-3 was almost 3000-fold higher (15,059 pg/mg), and expression of MMP-8 was 500-fold higher (12,923 pg/mg) at 8 weeks post stroke in the area of liquefactive necrosis compared to naïve brain tissue.

MMP-2 is a gelatinase, MMP-3 is a stromelysin, and MMP-8 is a collagenase (Amin et al., 2016), and such proteases digest proteins to amino acids. The chronically high expression of MMPs is a probable mechanism through which the solid tissue of the brain liquefies and remains liquefactive for months following stroke. Further, this may be an explanation for the much later collagen deposition that occurs in the brain compared to the heart.

MMPs can degrade myelin and so the initial high production of MMP-2, MMP-3, and MMP-8 at 24 hours following stroke may occur as a microglia/macrophage response to damaged myelin (Chandler et al., 1995; D’Souza et al., 2002). However, the overproduction of MMPs in the central nervous system is neurotoxic due to their capacity to disrupt tight junctions and degrade intact myelin on healthy neurons. Furthermore, MMP-2 converts stromal cell derived factor 1 (SDF-1), which is produced by reactive astrocytes, into a highly neurotoxic fragment (Zhang et al., 2003; Fujimoto et al., 2008; Mroczko et al., 2013). Other degradative enzymes are also likely to be substantially and chronically elevated in areas of liquefactive necrosis following stroke. The full characterization of this repertoire is likely to reveal additional targets for reducing secondary neurodegeneration.

In this study, we discovered that liquefactive necrosis is resolved by 24 weeks in young adult C57BL/6 mice, however, in light of Cantuti-Castelvetri et al.’s recent finding that phagocytes from aged animals are less able to process myelin debris than young animals, aged mice may take considerably longer (Cantuti-Castelvetri et al., 2018). How long it takes for liquefactive necrosis to resolve in humans is still unknown. It is probable that it is influenced by age, comorbidities, genetics, infarct size, and stroke location, and it is also likely that it is causing different amounts of secondary neurodegeneration in different patients. This is an important area for future stroke research.

Further emphasizing the need for such research, although it has yet to be fully characterized the extent to which humans form cholesterol crystals in the brain following stroke, in 1987 Miklossy and Van der Loos reported that crystals of cholesterol esters are present for at least 20 months in the human brain after stroke, both within the area of infarction, as well as in the tracts originating in and passing through the region of infarction (Miklossy and Van der Loos, 1987; Miklossy et al., 1991).

The cytokine profile in the area of liquefactive necrosis resembles the cytokine profile present in atheroma (Braunersreuther et al., 2007). For example, there is the chronic presence of MCP-1, IP-10, and RANTES, known participants in atheroma. Our previous data demonstrate that the human brain at the stage of liquefactive necrosis also contains many of the cytokines found in atheroma including MCP-1, IP-10, MIP-1α, and MIP-1β, (Nguyen et al., 2016). The presence of these cytokines not only demonstrates overlap between the molecular profile of atherosclerosis and liquefactive necrosis, but it also provides additional targets for intervention. For instance, blocking MCP-1, IP-10, and RANTES results in atherosclerosis regression in mice (Braunersreuther et al., 2007). With regard to the relative abundance of the cytokines present in the area of liquefactive necrosis, of the cytokines we analyzed, the most abundant cytokine was OPN. OPN is also one of the most abundant cytokines expressed in atheroma (Scatena et al., 2007).

There is a direct link between OPN and MMP expression. Extracellular MMP inducer (EMMPRIN) is an upstream inducer of several MMPs and is a master regulator of MMP production (Agrawal and Yong, 2011). The COOH-terminus of OPN, which is released upon thrombin cleavage and contains a cryptic SLAYGLR motif, is an EMMPRIN agonist. Specifically, the thrombin cleaved COOH-terminal fragment of OPN forms a complex with cyclophilin C, which upon engagement of EMMPRIN has been shown in murine mammary epithelial tumor cell lines to lead to phosphorylation of AKT and MMP-2 activation and secretion (Mi et al., 2007). Cyclophilin C and cyclophilin-C-associated protein (CyCap) are expressed by macrophages in response to tissue damage (Kong et al., 2007), and in the brain, cyclophilin C complexes with CyCap to activate microglia via the calcineurin/NFAT ((nuclear factor of activated T-cells) pathway (Yamaguchi et al., 2011). This may explain why MMP-2 and MMP-8 levels were substantially reduced in the infarct in the OPN deficient mice.

Further supporting a critical role of OPN in the mechanism of liquefactive necrosis, OPN is important in promoting the retention of macrophages at sites of chronic inflammation (Scatena et al., 2007), and is known to regulate chronic inflammatory responses to foreign bodies such as cholesterol crystals. This is evidenced by the fact that renal epithelial cells increase the production of OPN in response to exposure to calcium oxalate crystals (Khan et al., 2002), there are deficits in macrophage accumulation in OPN^-/-^ mice in models of biomaterial implantation (Scatena et al., 2007), and Oh and colleagues recently demonstrated that the thrombin cleaved COOH-terminal fragment of OPN plays a role in multinucleated giant cell formation (Oh et al., 2014).

With regard to therapeutic intervention, G-CSF, IL-1β, IP-10, MCP-1, MIP-1α, MIP-1β, RANTES, TNF-α, MMP-2, and MMP-8 were substantially reduced in OPN^-/-^ mice 7 weeks following stroke, and these mice exhibited less secondary neurodegeneration in the peri-infarct location. Although we do not know at this time if reduced secondary neurodegeneration in OPN^-/-^ mice is due to enhanced neurovascular repair, or rather due to the inflammatory milieu in the area of liquefactive necrosis being less toxic to surrounding cells, these possibilities are not mutually exclusive. Although the administration of OPN acutely is neuroprotective (Meller et al., 2005; Doyle et al., 2008), these data suggest that reducing OPN activity in the chronic inflammatory process may be a target for therapeutic intervention in recovering stroke patients. In support of this, it has been shown that OPN deficiency attenuates MMP activity and plaque accumulation in mouse models of atherosclerosis (Bruemmer et al., 2003; Matsui et al., 2003).

However, tempering the potential of targeting OPN expression as a means of improving stroke recovery in humans, the OPN^-/-^ mice in this study exhibited a significant increase in the amount of cholesterol crystals present in the area of liquefaction compared to their wildtype counterparts. Our interpretation of this finding is that because OPN appears to be a key driver of the chronic inflammatory response to stroke, the absence of OPN may be slowing down the clearance of myelin debris, thereby enabling more cholesterol crystals to form. This adds a nuance to our data as it indicates that targeting OPN to reduce chronic inflammation in stroke recovery comes at the cost of additional cholesterol crystal formation. Future studies are necessary to address the translational potential of targeting OPN following stroke with regard to whether the absence of OPN and the formation of additional cholesterol crystals actually leads to a worse overall outcome at time points later than 7 weeks.

Surprisingly, genetically ablating the NLRP3 inflammasome, which is activated by intracellular cholesterol crystals, had little impact on cytokine production or the MMP response to stroke. Although IL-1 levels were reduced in the area of liquefaction as expected, no other proteins measured were significantly altered. This unexpected result suggests that the NLRP3 inflammasome is less integral to the pathophysiology of liquefactive necrosis than OPN. An explanation for this result is that there are consequences of myelin debris clearance in addition to the formation of cholesterol crystals that are preventing the inflammatory response to stroke from resolving as efficiently as the inflammatory response to MI. Examples are the formation of oxidized LDL and oxysterols. Oxidized LDL can bind to pattern recognition receptors, including TLRs, and can thereby directly trigger pro-inflammatory signaling pathways (Remmerie and Scott, 2018). Oxysterols are a product of cholesterol processing in foam cells, and they are another important driver of inflammation in atherosclerotic lesions (Gargiulo et al., 2011; Poli et al., 2013). More research is needed to determine the relative contributions of cholesterol crystals, oxidized LDL, oxysterols, and other myelin catabolites to the pathogenicity of chronic stroke infarcts.

CD36 is a key regulator of lipid droplet formation in atherogenesis due to its ability to bind to lipoproteins, oxidized phospholipids, fatty acids, and oxidized LDL. However, in this study the genetic ablation of CD36 had little impact on chronic cytokine production and MMP expression after stroke. This lack of an effect may be due to redundancy in scavenger receptor expression on the surface of microglia and macrophages.

In conclusion, myelin has evolved to form a stable and long-lived insulating sheath that is difficult to degrade (Schmitt et al., 2015). Consequently, following a sizable CNS insult such as a stroke, the production of myelin debris and the release of the associated cholesterol may be overwhelming the cholesterol degradation capacities of phagocytosing microglia and macrophages. Thus, treatments that target cholesterol overloading within microglia and macrophages, and the sustained production of degradative enzymes, may help to prevent the damaging effects of liquefactive necrosis following stroke.

## Ethical approval

All applicable international, national, and/or institutional guidelines for the care and use of animals were followed. As such, all experiments were conducted in accordance with protocols approved by the Animal Care and Use Committee of the University of Arizona and were performed based on the NIH Guide for the Care and Use of Laboratory Animals.

## Acknowledgements

None

